# SARS-CoV-2 induces human plasmacytoid pre-dendritic cell diversification via UNC93B and IRAK4

**DOI:** 10.1101/2020.07.10.197343

**Authors:** Fanny Onodi, Lucie Bonnet-Madin, Laurent Meertens, Léa Karpf, Justine Poirot, Shen-Ying Zhang, Capucine Picard, Anne Puel, Emmanuelle Jouanguy, Qian Zhang, Jérôme Le Goff, Jean-Michel Molina, Constance Delaugerre, Jean-Laurent Casanova, Ali Amara, Vassili Soumelis

## Abstract

Several studies have analyzed antiviral immune pathways in late-stage severe COVID-19. However, the initial steps of SARS-CoV-2 antiviral immunity are poorly understood. Here, we have isolated primary SARS-CoV-2 viral strains, and studied their interaction with human plasmacytoid pre-dendritic cells (pDC), a key player in antiviral immunity. We show that pDC are not productively infected by SARS-CoV-2. However, they efficiently diversified into activated P1-, P2-, and P3-pDC effector subsets in response to viral stimulation. They expressed CD80, CD86, CCR7, and OX40 ligand at levels similar to influenza virus-induced activation. They rapidly produced high levels of interferon-α, interferon-λ1, IL-6, IP-10, and IL-8. All major aspects of SARS-CoV-2-induced pDC activation were inhibited by hydroxychloroquine. Mechanistically, SARS-CoV-2-induced pDC activation critically depended on IRAK4 and UNC93B1, as established using pDC from genetically deficient patients. Overall, our data indicate that human pDC are efficiently activated by SARS-CoV-2 particles and may thus contribute to type I IFN-dependent immunity against SARS-CoV-2 infection.

## Introduction

Severe Acute Respiratory Syndrome-Coronavirus-2 (SARS-CoV-2) is the third zoonotic coronavirus that emerged in the last two decades. SARS-CoV-2 is the causative agent of coronavirus disease 2019 (COVID-19) that appeared in late 2019 in Wuhan, Hubei province in China (Nandakumar, 2020; Tay et al., 2020). SARS-CoV-2 became rapidly pandemic, and infection have now been detected in 216 countries and territories, and is responsible for approximately 10 million confirmed cases and > 1 million deaths (WHO, weekly epidemiological update - 24 November 2020). SARS-CoV-2 infection may lead to a diversity of clinical presentations, ranging from asymptomatic or mild “flu-like” syndrome in >98% of patients, to severe and sometimes life-threatening acute respiratory failure in <2% of infected individuals (Tang et al., 2020). Disease aggravation usually occurs after 8 to 10 days following initial symptoms (Tang et al., 2020). At this late stage, three main factors were shown to correlate with the progression and severity of the infection (Tang et al., 2020): 1) viral persistence was evidenced in the lung and systemic circulation, although it is not constant (Tang et al., 2020), 2) an excess production of pro-inflammatory cytokines, such as IL-1β and IL-6 (Tay et al., 2020), 3) a defect in type I interferon (IFN) production, especially in critically ill patients (Tay et al., 2020; Acharya et al., 2020). Whether an imbalance between inflammatory cytokines and type I IFN occurs early in the disease, at the stage of the primary infection, and whether the virus itself may be responsible is currently unknown. More recent studies clarified the pathogenesis of life-threatening COVID-19 pneumonia in about 15% of patients. While some patients carry inborn errors of TLR3-and IRF7-dependent type I IFN production and amplification (Zhang et al., 2020b), at least 10% of patients have pre-existing neutralizing auto-Abs to type I IFNs (Bastard et al., 2020). These studies suggested that the pathogenesis of life-threatening COVID-19 proceeds in two steps, with defective type I IFNs in the first days of infection unleashing excessive inflammation from the second week onward (Zhang et al., 2020a).

Investigating the early innate immune response to SARS-CoV-2 is essential both to understand the mechanisms underlying an efficient viral control, and to shed light on possible subsequent life-threatening complications. Plasmacytoid pre-dendritic cells (pDC) play a particularly important role as the major source of type I IFN in response to viral infection, due to their constitutively high levels of IRF7 expression (Liu, 2005). PDC can sense a large array of viruses including the coronaviruses SARS-CoV, murine hepatitis virus (MHV) and the middle east respiratory syndrome coronavirus (MERS) (Cervantes-Barragan et al., 2007, 2012; Raj et al., 2014; Scheuplein et al., 2015). PDC recognize MERS through TLR-7 (Scheuplein et al., J Virol 2015). They respond to viruses by producing innate cytokines, including all forms of type I IFNs (α and β), type III IFN, and inflammatory cytokines, in particular TNF-α and IL-6 (Liu, 2005; Gilliet et al., 2008). However, different viruses may induce different cytokine patterns (Thomas et al., 2014), possibly creating an imbalance between IFN versus inflammatory cytokine response. Additionally, some viruses were shown to subvert pDC functions through different mechanisms not necessarily related to productive infection. This is the case for human immunodeficiency virus, which may induce pDC apoptosis in vitro (Meyers et al., 2007) and pDC depletion in vivo (Soumelis et al., 2001; Meera et al., 2010). Human hepatitis C virus can inhibit IFN-α production by pDC through the glycoprotein E2 binding to BDCA-2 (Florentin et al., 2012). Human papillomavirus induces very low IFN response in pDC (Bontkes et al., 2005), which may be due to impaired Toll-Like Receptor (TLR) −7 and −9 signaling (Hirsch et al., 2010). Whether SARS-CoV-2 induces efficient pDC activation, or may interfere with various biological pathways in pDC is currently unknown.

## Results and Discussion

### SARS-CoV-2 induces activation and diversification of primary human pDC

In order to efficiently recapitulate SARS-CoV-2-pDC interactions, we used two strains of SARS-CoV-2 primary isolates. Their viral genome sequences were nearly identical with 99,98% identity. Sequence comparison with reference strain Wuhan-Hu-1 (NCBI accession number NC_045512.2) showed that both strains contain a subset of mutations (C241T; C30307T; A23403G and G25563T), characteristic of the GH clade based on GISAID nomenclature. Human primary pDC were purified from healthy donor (HD) peripheral blood mononuclear cells (PBMC) by cell sorting. First, we asked whether SARS-CoV-2 was able to induce pDC activation, and diversification into IFN-producing and/or T cell stimulating effectors, as we previously described for influenza virus A (Flu) (Alculumbre et al., 2018). After 24 hours of culture, SARS-CoV-2-activated pDC efficiently diversified into P1 (PD-L1^+^CD80^-^), P2 (PD-L1^+^CD80^+^), and P3 (PD-L1^+^CD80^+^) pDC subsets, similar to Flu stimulation (Fig. 1A). P1-, P2-, and P3-pDC were all significantly induced by SARS-CoV-2 and Flu, as compared to medium control (Fig. 1 B). In parallel, we observed a sharp decrease in non-activated P0-pDC (PD-L1^-^CD80^-^) (Fig. 1 A and B). SARS-CoV-2-induced pDC activation was comparable with magnetically-versus FACS-sorted pDC (Fig. S1 A and B), confirming that both methods are suitable for subsequent experiments. All main findings were confirmed on at least three independent experiments using FACS-sorted pDC, with a protocol that excluded AS-DC, a rare dendritic cell (DC) subset that shares some markers and functional features with pDC (Villani et al., 2017), based on CD2, CD5 and AXL expression (Fig. S1 A). PDC activation and diversification was observed with two independent primary SARS-CoV-2 strains (Fig. 1 C), which both induced similar proportions of P1-P3 subsets. PDC diversification was also observed by co-culturing of pDC with SARS-CoV-2-infected Vero E6 cells with a similar efficiency than free SARS-CoV-2 (Fig. S1 C). SARS-CoV-2 improved pDC viability as compared to medium condition (Fig. 1 D), which is compatible with subsequent effector functions.

**Figure 1.**
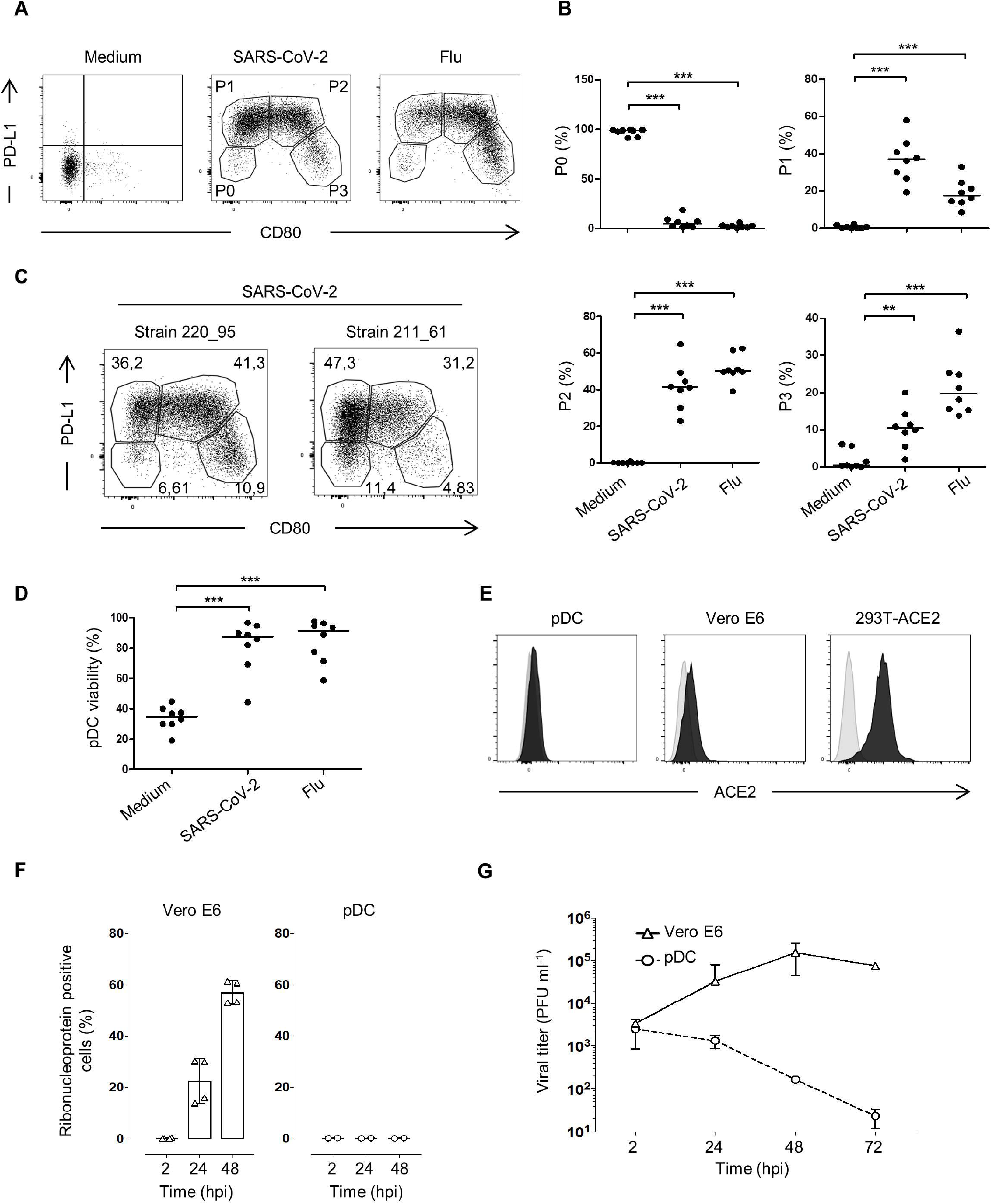
SARS-CoV-2 induces activation and diversification of primary human pDC. Sorted blood pDC from healthy donors were cultured for 24h with either Medium, SARS-CoV-2, or Influenza virus A (Flu). **(A)** Dotplot showing pDC activation and diversification through the expression of PD-L1 and CD80 into P1-, P2-, and P3-subpopulations. Results from one healthy donor representative of n=8. **(B)** Quantification of the three populations. Bars represent medians of n=8 healthy donors from 6 independent experiments. **P□< 0.01; ***P□< 0.005 Mann Whitney test. **(C)** Dotplot showing pDC activation from different strains of SARS-CoV-2 isolated from two patients. Results from one healthy donor representative of n=3. **(D)** Percentage of live pDC after 24h of culture with either Medium, SARS-CoV-2, or Influenza virus A (Flu). n=8 healthy donors from 6 independent experiments. ***P□< 0.005 Mann Whitney test. **(E)** Histogram of ACE2-expression on pDC, Vero E6 and 293T-ACE2 (black) compared to the isotype (light grey). Results from one experiment representative of n=3. **(F)** Intracellular production of SARS-CoV-2 Ribonucleoprotein in Vero E6 and pDC at 2, 24 or 48h post infection (hpi) with SARS-CoV-2. Results from one experiment representative of n=3. **(G)** Infectious viral titers in the supernatants of SARS-CoV-2-infected Vero E6 and pDC at 2, 24, 48 or 72h post infection (hpi). Results from one experiment representative of n=3.

The fact that SARS-CoV-2 induced a complete diversification into the three activated pDC subsets suggests that different aspects of pDC-mediated immunity are engaged following viral challenge: type I interferon production and innate immunity mediated by P1-pDC, T cell stimulation mediated by P3-pDC, and a mixed innate and adaptive effector function through P2-pDC (Alculumbre et al., 2018). Such a strong innate immune response could be central to the systemic and local inflammatory manifestations in primary symptomatic SARS-CoV-2 Infection (Asselah et al., 2020).

### Human pDC are not productively infected by SARS-CoV-2

Next, we asked whether SARS-CoV-2 -induced pDC activation was dependent on productive infection. We first checked whether pDC express at their cell surface ACE2, the major SARS-CoV-2 entry receptor (Hoffmann et al., 2020). No significant expression was detected using an anti-ACE2-specific antibody, as compared to a low and high expression on Vero E6 and 293T-ACE2 cell lines, respectively (Fig. 1 E). The ability of pDC to replicate SARS-CoV-2 was then assessed. Human pDC were challenged with SARS-CoV-2 strain 220_95 at MOI of 2, and cultured for 2h, 24h or 48 h. Our results showed that pDC were refractory to SARS-CoV-2 infection, as evaluated by quantifying 1) the intracellular production of the nucleoprotein antigen (N) (Fig. 1 F), or the accumulation of viral RNA in SARS-CoV-2-infected cells (Fig. S1 D), and 2) the release of infectious progeny virus in the supernatants of infected cells using plaque assays (Fig. 1 G). As positive control, the permissive Vero E6 cells produced high level of the N antigen, increased viral RNA overtime (Fig. S1 D), and high viral titers following SARS-CoV-2 incubation (Fig. 1 G). Similar results were obtained with pDC isolated from three independent donors (Fig. S1 E). Overall, these results show that pDC are resistant to SARS-CoV-2 infection, and are efficiently activated by the virus independently of ACE2 expression. Their viability was not affected by SARS-CoV-2 challenge.

### Upregulation of major immune checkpoints on SARS-CoV-2-activated pDC

Activating immune checkpoints play a key role in T cell stimulation, and serve as surrogate markers of DC differentiation (Guermonprez et al., 2002). We first assessed the dose-dependent effect of SARS-CoV-2 on CD80 expression and subset diversification. CD80 was induced in a dose-dependent manner by SARS-CoV-2 at MOI 0.04 to 1 (Fig. 2 A). This was accompanied by an increase in P3-pDC subset, and a slight decrease in P1-pDC (Fig. 2 B). A detailed phenotypic analysis was subsequently performed on pDC after 24 and 48 hours of culture with SARS-CoV-2 (Fig. 2 C and Fig. S2 A). Diversification was observed at both time points, with a slight increase in P3-pDC at 48 hours (Fig. S2 A). P2- and P3-pDC significantly upregulated CD80, CD86, CCR7, and OX40L, as compared to non-activated P0-pDC, in both SARS-CoV-2 and Flu conditions (Fig. 2 C). PD-L1, and CD62L, an integrin that promotes lymph node homing, were both higher on P1- and P2-pDC (Fig. 2 C). Expression of checkpoint molecules persisted at 48h, especially the higher CD80 and CD86 expression on P3-pDC (Fig. S2 B).

**Figure 2.**
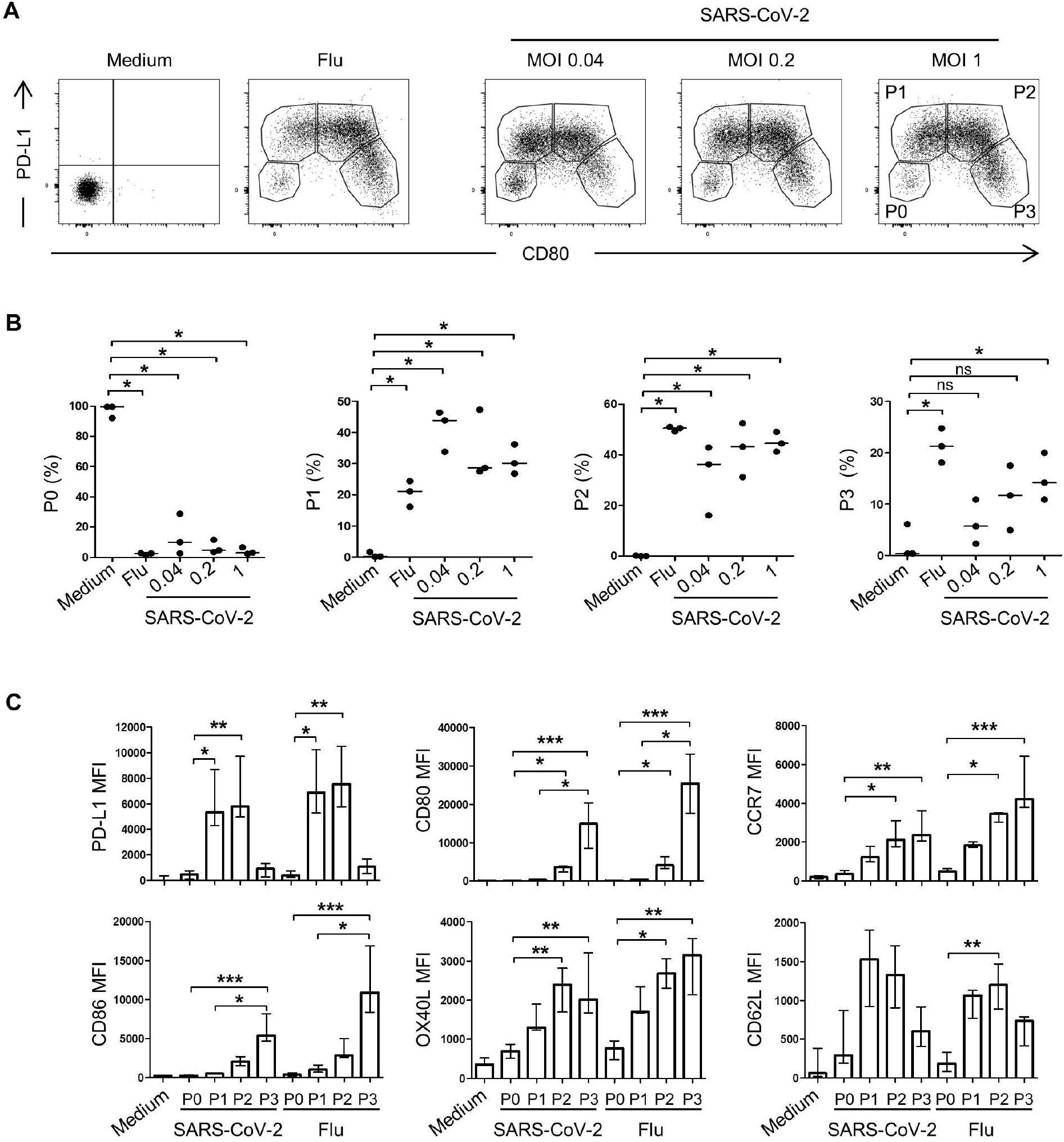
SARS-CoV-2 induces pDC activation in a dose dependent manner. Sorted blood pDC from healthy donors were cultured for 24h with either Medium, Influenza virus A (Flu), or SARS-CoV-2 at a MOI of 0.04, 0.2, or 1. **(A)** Dotplot showing pDC activation through the expression of PD-L1 and CD80. Results from one healthy donor representative of n=3. **(B)** Quantification of the three populations. Bars represent medians of n=3 healthy donors from 3 independent experiments. *P□< 0.05; ns: not significant; Mann Whitney test. **(C)** PDC geometric mean (MFI) of activation markers after 24h of culture with either Medium, Influenza virus A, or SARS-CoV-2 at a MOI of 1. Histograms represent medians and bars interquartile of n=5 healthy donors from 3 independent experiments. *P□< 0.05; **P□< 0.01; ***P□< 0.001; Kruskal-Wallis with Dunn’s multiple comparison’s post test.

### Efficient production of type I and type III interferons by SARS-CoV-2-activated pDC

A key and defining function of pDC is their ability to produce large amounts of type I IFN, driven by their constitutively high levels of IRF7 (Liu, 2005). We measured the production of several cytokines at the protein level after 24 hours of culture. Both SARS-CoV-2 and Flu induced high levels of IFN-α2 and IFN-λ1, both being critical anti-viral effector cytokines (Fig. 3 A). IFN-α2 levels following SARS-CoV-2 activation reached up to 80 ng/ml, indicating a very efficient activation. The chemokine IP-10 was also significantly induced (Fig. 3 A), possibly due to an autocrine IFN loop (Blackwell and Krieg, 2003). Inflammatory cytokines IL-6 and IL-8 were comparably induced by SARS-CoV-2 and Flu (Fig. 3 A). However, TNF-α levels were marginally induced by SARS-CoV-2, when compared with Flu activation (Fig. 3 A). Cytokine production was maintained after 48 hours of viral activation (Fig. S2 C). Secreted protein levels were similar to 24h levels for most cytokines. Interestingly, IFN-α levels rose by 3-fold between 24h and 48h for one donor (Fig. 3 A and S2 C), indicating the possibility of increased production. Such strong IFN producer suggests a potential virus controller.

**Figure 3.**
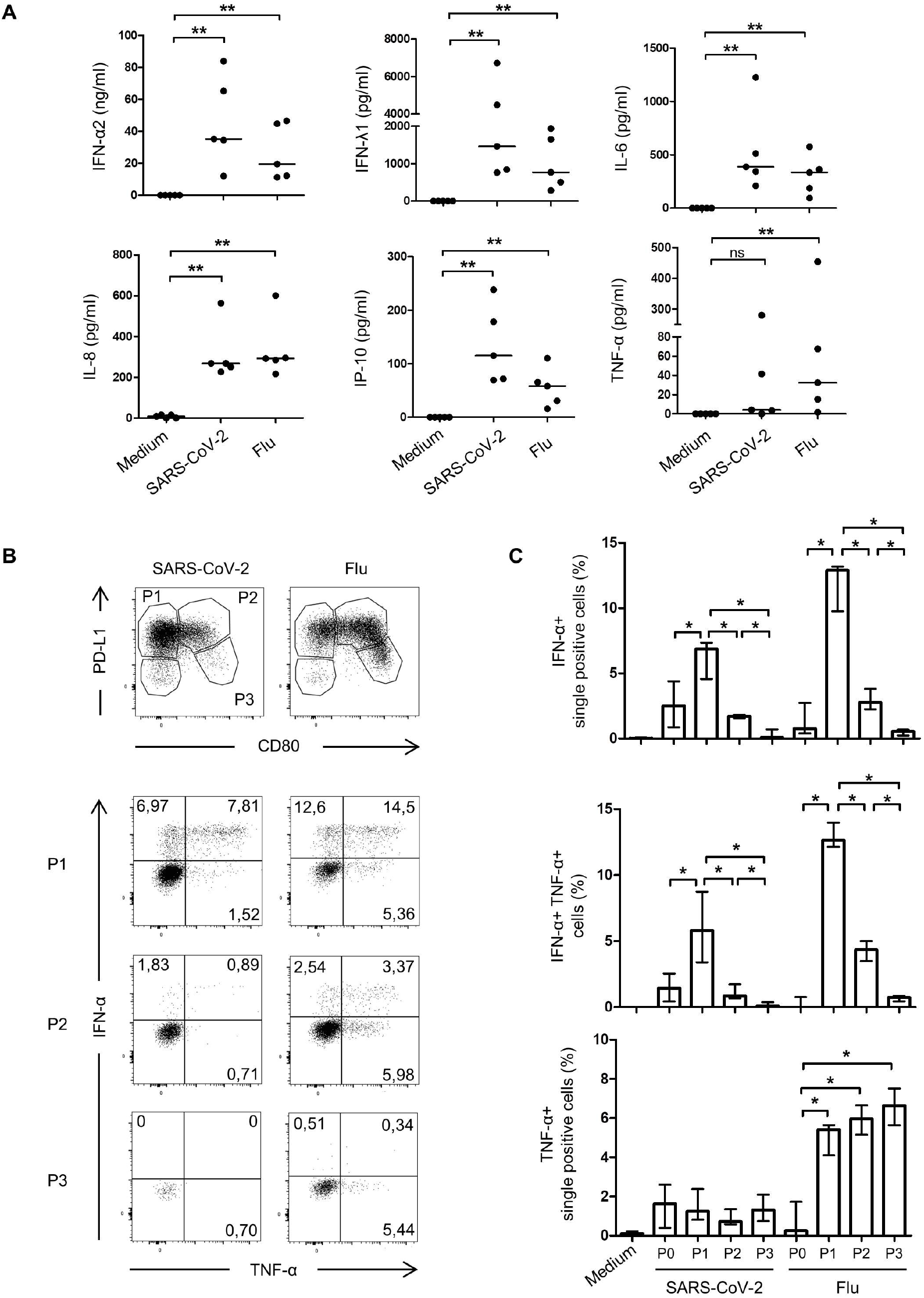
SARS-CoV-2-activated pDC produce pro-inflammatory cytokines. Sorted blood pDC from healthy donors were cultured for 24h with either Medium, Influenza virus A (Flu), or SARS-CoV-2. **(A)** Quantification of secreted pro-inflammatory cytokines after 24h of culture. Bars represent medians of n=5 healthy donors from 3 independent experiments. **P□< 0.01; ns: not significant; Mann Whitney test. **(B)** Dotplot showing pDC activation through the expression of PD-L1 and CD80 (upper plots), and intracellular IFN-α and TNF-α in P1, P2 or P3 populations (lower plots). Results from one healthy donor representative of n=4. **(C)** Percentages of IFN-α single positive, IFN-α^+^ TNF-α^+^ double positive, and TNF-α single positive cells in P0, P1, P2 or P3 populations. Histograms represent medians and bars interquartile of n=4 healthy donors from 3 independent experiments. *P□< 0.05; Mann Whitney test.

We have shown that P1-, P2, and P3-pDC exhibit different functions in response to Flu activation, including different cytokine profiles (Alculumbre et al., 2018). We questioned whether this functional specialization was similar following SARS-CoV-2 challenge. PDC were cultured with either Flu or SARS-CoV-2 for 24h, and P1-, P2- and P3-pDC differentiation was assessed, along with intracellular IFN-α and TNF-α staining (Fig. 3 B). In the SARS-CoV-2 condition, IFN-α was essentially produced by P1-, and to a lower extent by P2-pDC, with 12% and 3% of IFN-α positive cells, respectively. TNF-α was also mainly produced by P1-pDC, with 7% expressing-cells (Fig. 3 B and C). P3-pDC produced very little cytokines. Similar results were observed following Flu challenge (Fig. 3 C). These data showed that IFN-α and TNF-α production are features of P1-pDC, as well as P2-pDC, and that SARS-CoV-2 induces a functional specialization of pDC subsets, similar to Flu.

Because the oropharyngeal mucosa is an entry site for SARS-CoV-2, we aimed at validating our results using pDC purified from tonsils. SARS-CoV-2 induced a marked diversification of tonsillar pDC into all three activated subsets (Fig. S2 D). Tonsillar pDC efficiently produced IFN and inflammatory cytokines in response to SARS-CoV-2 (Fig. S2 E). Overall, our results establish SARS-CoV-2 as a very efficient inducer of type I and type III IFN responses. Inflammatory cytokines were induced at similar level than Flu activation, without any significant imbalance that would be suggestive of excessive inflammatory response.

### SARS-CoV-2-induced pDC activation is inhibited by hydroxychloroquine

We then asked whether pharmacological agents could modulate pDC diversification and cytokine production. Hydroxychloroquine (HCQ) is known to inhibit endosomal acidification which may diminish pDC activation (Sacre et al., 2012). Additionally, it is being tested in several clinical studies as a potential treatment for COVID-19 (Das et al., 2020). Hence, we addressed its role in SARS-CoV-2-induced pDC activation. Following 24 hours of culture, we found that HCQ inhibited pDC diversification in response to SARS-CoV-2, which is similar to the decrease observed with Flu, used as a positive control (Fig. 4 A). In particular, P2- and P3-pDC differentiations were almost completely inhibited (Fig. 4 B). Inhibition of SARS-CoV-2-induced pDC diversification by HCQ was dose-dependent (Fig. S3 A). The significant decrease in P3-pDC was paralleled by a decrease in CD80, CD86, and CCR7 expression (Fig. 4 C and D). OX40-ligand expression was not significantly affected by HCQ (Fig. 4 C and D). However, HCQ inhibited the appearance of an OX40-ligand^high^ pDC population (Fig. S3 B and C), which may impact subsequent T cell activation. Last, we assessed the effect of HCQ on innate pDC functions. We measured cytokine production after 24 hours of SARS-CoV-2-induced pDC activation in the presence or absence of HCQ. We found that IFN-α and λ levels were decreased by HCQ (Fig. 4 E). This was also the case for IL-6 and IL-8, with a much lesser impact on IP-10 (Fig. 4 E). Together, these results show that HCQ inhibits SARS-CoV-2-induced pDC diversification and cytokine production.

**Figure 4.**
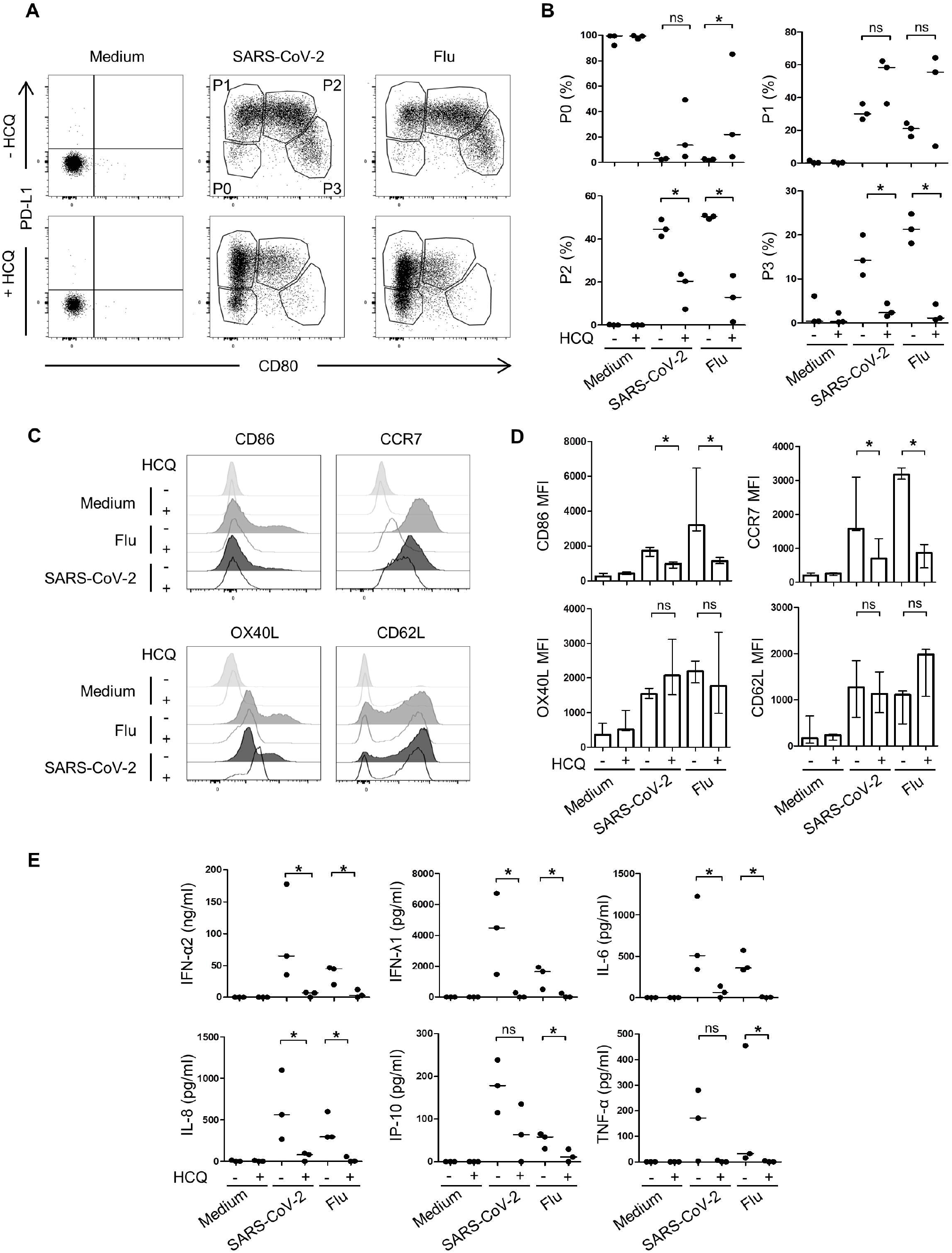
SARS-CoV-2-induced pDC activation is inhibited by hydroxychloroquine. Sorted blood pDC from healthy donors were cultured for 24h with either Medium, Influenza virus A (Flu), or SARS-CoV-2 at a MOI 1 with or without the presence of hydroxychloroquine (HCQ). **(A)** Dotplot showing pDC diversification in P1-, P2-, and P3-subpopulations in the presence of HCQ. Results from one healthy donor representative of n=3. **(B)** Quantification of the three populations. Bars represent medians of n=3 healthy donors from 3 independent experiments. *P□< 0.05; ns: not significant; Mann Whitney test. **(C)** Histograms of pDC’s activation markers. Results from one healthy donor representative of n=3. **(D)** Geometric mean (MFI) of activation markers. Histograms represent medians and bars interquartile of n=3 healthy donors from 3 independent experiments. **(E)** Quantification of pro-inflammatory cytokines production. Bars represent medians of n=3 healthy donors from 3 independent experiments. *P□<c0.05; ns: not significant; Mann Whitney test.

### pDC response to SARS-CoV-2 depends on IRAK4 and UNC93B1

In order to gain mechanistic insights, we took advantage of studying pDC from patients with genetic deficiencies in the TLRs signaling pathway (Rothenfusser et al., 2002; Casanova et al., 2011). While pDCs do not express detectable TLR3, they express TLR7 and TLR9 (Liu, 2005). We investigated pDC from patients homozygous for germline mutations, resulting in loss of function of IRAK4 (one patient), which with MyD88 is required for signaling of IL-1Rs and TLRs other than TLR3 (Frazão et al., 2013), or UNC93B1 (two patients), which is required for signaling of endosomal TLR3, TLR7, TLR8, and TLR9 (Picard et al., 2003; Casrouge et al., 2006; Lee and Barton, 2014). PDC from one TLR3 deficient patient was used as negative control (Zhang et al., 2007; Guo et al., 2011) (Fig. 5). PDC from IRAK4- and UNC93B1-deficient patients did not diversify into P1, P2 and P3 activated subpopulation, when compared with HD (Fig. 5 A). This lack of diversification was associated with a complete absence of IFN-α2, IP-10, and IL-6 secretion, establishing a critical role of these molecular nodes in SARS-CoV-2 activation of pDC (Fig. 5 B). On the contrary, TLR3 deficiency did not significantly impact SARS-CoV-2-induced pDC activation, associated with diversification, and secretion of anti-viral cytokines (Fig. 5 A and B). Taken together, these results identify IRAK4 and UNC93B1 as two critical players in controlling SARS-CoV-2 pDC activation.

**Figure 5.**
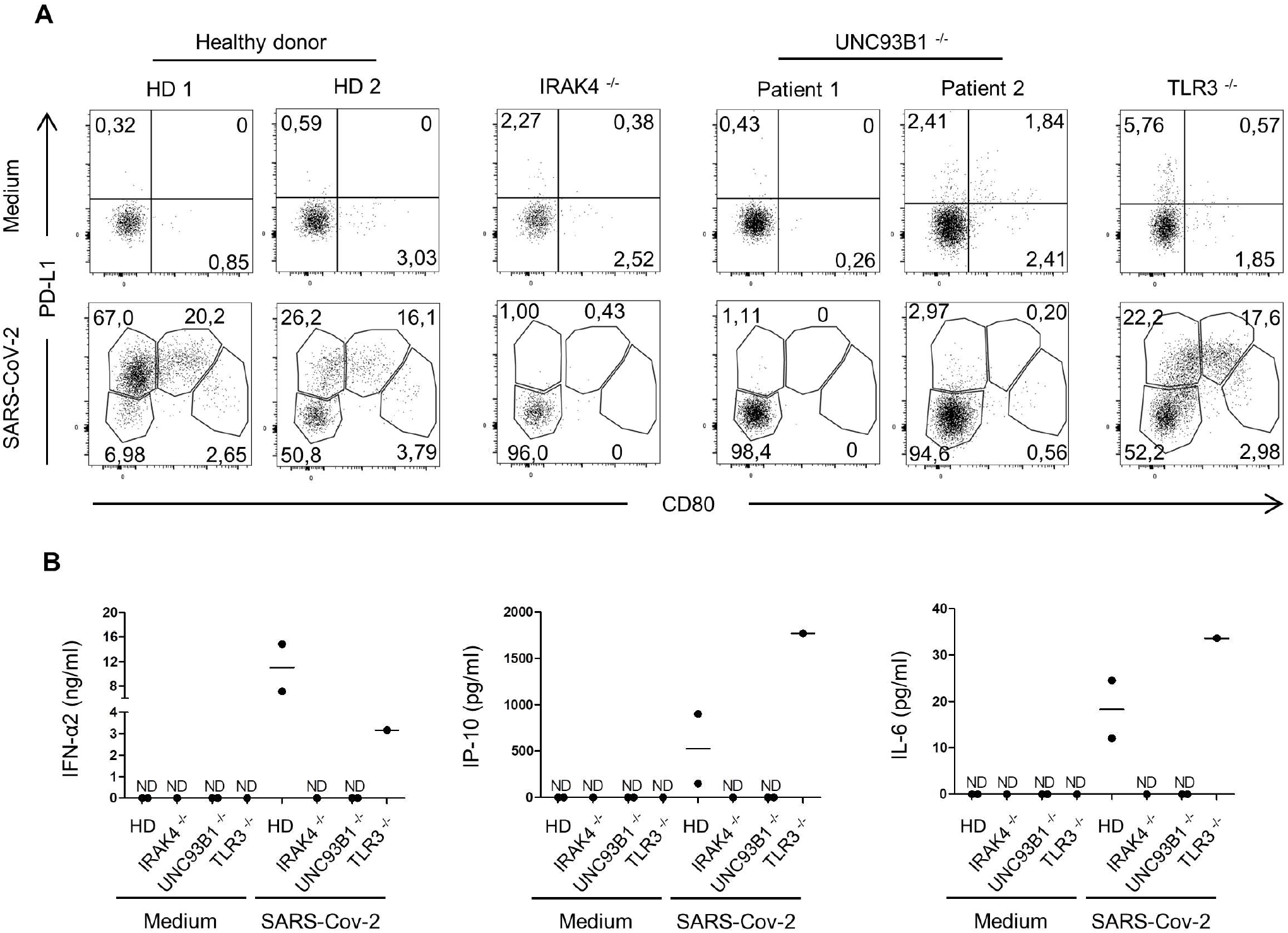
SARS-CoV-2-induced pDC activation requires IRAK4 and UNC93B1. Sorted blood pDC from mutated patients and healthy donors were cultured for 24h with either Medium or SARS-CoV-2. **(A)** Dotplot showing pDC diversification in P1-, P2-, and P3-subpopulations from magnetically sorted blood pDC from homozygous IRAK4^-/-^ (n = 1), UNC93B1^-/-^ (n = 2), TLR3^-/-^ (n = 1) donors, and gender aged-matched healthy donors (n = 2) were cultured for 24h with either Medium, or SARS-CoV-2 at a MOI 1. **(B)** Quantification of pro-inflammatory cytokines production by in the supernatant of activated pDC for 24h in response to SARS-CoV-2 challenge. Bars represent medians; ND: Not Detectable.

In this study, we have analyzed the interaction between primary SARS-CoV-2 isolates and human primary pDC. Our results demonstrate that SARS-CoV-2 is a strong IFN inducer by efficiently stimulating primary pDC. Both type I and type III IFNs were induced at high levels in P1-pDC upon SARS-CoV-2 stimulation. This strongly suggests that the defects observed in critically ill COVID-19 patients are acquired during disease evolution through secondary events, not necessarily associated to direct effect of the virus. Possible mechanisms could be related to inflammatory cytokines, such as TNF, and endogenous glucocorticoid response, both being able to promote pDC apoptosis (Abe and Thomson, 2006; Lepelletier et al., 2010).

Our results are consistent with those obtained using other coronaviruses, such as SARS-CoV (Cervantes-Barragan et al., 2007), MERS (Raj et al., 2014), and MHV (Cervantes-Barragan et al., 2007, 2012), which all induced an efficient IFN response by pDC independently of viral replication. However, pDC response to MERS depended on binding to its receptor DPP4 (Raj et al., 2014). In our study, viral sensing was independent of the expression of the ACE2 entry receptor. Importantly, pDC response to SARS-CoV-2 was dependent on two critical molecular components downstream of TLRs, IRAK4 and UNC93B1. These results are consistent with studies showing the importance of IRAK4 in pDC response to immune complexes through TLR7 (Hjorton et al., 2018, 4; Corzo et al., 2020). UNC93B1 was reported to regulate transport, stability, and function of endosomal TLR3, TLR7, TLR8, and TLR9 (Lee and Barton, 2014). Our studies are also consistent with our previous studies of pDCs in patients with inherited IRF7 deficiency (Ciancanelli et al., 2015; Zhang et al., 2020b). Indeed, we showed that IRF7-deficient pDCs did not produce type I IFNs other than IFN-β in response to influenza virus or SARS-CoV-2 (Zhang et al., 2020b). In our study, we provide direct evidence for a specific role of UNC93B1 in primary human pDC. The only shared features of human IRAK4 and UNC93B deficiencies being their control of cellular responses to TLR7, TLR8, and TLR9 (Casanova et al., 2011), we can infer from these studies that human pDC recognize SARS-CoV-2 via one or another of them, or a combination thereof. The abundant expression of TLR7 on human pDC suggests that this receptor may play an important role in this process (Gilliet et al., 2008; van der Made et al., 2020). It will be important to analyze the natural history of SARS-CoV-2 infection in patients with inborn errors of the TLR7, TLR8 and TLR9 pathway, including IRAK4, MyD88, and UNC93B deficiencies.

Type I IFNs are critical cytokines to control viral replication. Several chronic viral infections were associated with poor type I IFN responses (Snell et al., 2017; Marsili et al., 2012;). In COVID-19 patients, decreased serum levels of type I IFN were associated with severity in late stage infection, and increased viral load (Tay et al., 2020; Acharya et al., 2020). Defective type I IFN immunity was shown to underlie life-threatening COVID-19 pneumonia in either of two ways, via inborn errors of TLR3- and IRF7-dependent type I IFN immunity (Zhang et al., 2020b) or via pre-existing neutralizing auto-Abs to type I IFN (Bastard et al., 2020). This raised the question of the cellular basis of critical COVID-19 in patients with defective type I IFN immunity (Zhang et al., 2020a). Indeed, type I IFN is produced by all cells of the human body, including pulmonary epithelial cells, which also respond to type I IFNs, while it can be produced at much higher levels by pDC, which are the only cells expressing constitutively high levels of IRF7 (Liu, 2005).

The difficulty in designing current and future COVID-19 treatment strategies lies in part in the complexity of the host-virus interactions. Continued efforts in mapping and dissecting immune effector pathways to SARS-CoV-2 will be of major importance in order to design efficient treatment strategies adapted to each patient and stage of the infection.

## Materials and methods

### Patients

The experiments involving human subjects were conducted in accordance with local, national, and international regulations and were approved by the French Ethics Committee, the French National Agency for the Safety of Medicines and Health Products, and the French Ministry of Research (protocols C10-13 and C10-16). Informed consent was obtained from all patients included in this study.

The two patients with UNC93B1 deficiency suffered from two herpetic encephalitis episodes during childhood and are carrying homozygous mutations leading to a premature stop codon (1034del4) or an alternative splice event (781G>A) (Casrouge et al., 2006). The TLR3-deficient patient also displayed herpetic encephalitis, and is carrying compound heterozygous mutations (P554S and E746X) (Zhang et al., 2007; Guo et al., 2011). The IRAK4-deficient patient is a 5 years old female, who presented cellulitis to *Staphylococcus aureus* and a neutropenia from the age of 2 years and is carrying a homozygous stop-gain mutation (Q293X).

### PDC isolation and cell culture

Buffy coats from healthy human donors were obtained from Etablissement Français du Sang, Paris, Saint-Antoine Crozatier blood bank. PBMC were isolated through Ficoll density gradient centrifugation (Ficoll-Paque; GE Healthcare). PDC were isolated through a first step of pDC magnetic sorting (Human Plasmacytoid DC Enrichment Kit; StemCell), and subsequent flow cytometric sorting on the basis of live, lineage– (CD16, CD14, CD19, CD20, CD56 and CD3), CD11c– CD4+, and CD2– CD5– cells to a 98% purity. Due to some logistic issues, alternatively frozen PBMCs from Etablissement Français du Sang, Paris, Saint Louis blood bank, were thawed and placed at 37°C for 2h for cell recovery. PDC were then magnetically sorted (Human Plasmacytoid DC Enrichment Kit; StemCell). PDC enrichment was assessed based on the cytometric sorting panel, and was ranged from 71 to 90%. African green monkey kidney epithelial Vero E6 cells (ATCC, CRL-1586) were cultured in Dulbecco’s Modified Eagle Medium (DMEM; Thermo Fisher Scientific) supplemented with 10% FBS, 1% penicillin-streptomycin, 1% GlutaMAX and 25 mM Hepes

### SARS-CoV-2 primary strain isolation and amplification

SARS-CoV-2 viruses were isolated from nasopharyngeal swab specimens collected at Service de Virologie (Hospital Saint Louis, Paris). Samples were centrifugated at 4,000 x g for 10 min then filtered using a 0.45 μM filter, and diluted 1:1 with DMEM-4% (DMEM supplemented with 4% FBS, 1% penicillin-streptomycin, 1% GlutaMAX and 25 mM Hepes). Vero E6 cells were seeded in 96-well cell culture plate (15,000 cells/well), and incubated at 37°C with 200μl of inoculum and observed daily for cytopathogenic effects (CPE) by light microscopy. Substantial CPE were seen at 72-96 hours post inoculation. Culture supernatants were then collected, clarified by centrifugation, filtered using a 0.45 μM filter and kept at −80°C. We confirmed SARS-CoV-2 replication by RT-qPCR and whole viral genome sequences were obtained by next generation sequencing using Illumina MiSseq sequencers. Strains sequences have been deposited in the Global Initiative of Sharing All Influenza Data (GISSAID) database with accession ID EPI_ISL_469284 (220_95) and EPI_ISL_469283 (211_61). All viruses belong to the GH clade.

SARS-CoV-2 strains were further propagated on Vero E6 in DMEM-2% (DMEM supplemented with 2% FBS, 1% penicillin-streptomycin, 1% GlutaMAX and 25 mM Hepes). Viruses were passaged three times before being used for experiments. For the last passage, viruses were purified through a 20% sucrose cushion by ultracentrifugation at 80,000 x g for 2 hours at 4°C. Pellets were resuspended in HNE 1X pH7.4 (Hepes 5 mM, NaCl 150 mM, EDTA 0.1 mM), aliquoted and stored at −80°C.

Viruses titer was ascertained by plaque assays in Vero E6 cells and expressed as PFU per ml. Cells were incubated for 1 hour with 10-fold dilution of viral stocks. The inoculum was then replaced with Avicel 2.4% mixed at equal volume with DMEM supplemented with 4% FBS, 2% Glutamax, 50mM MgCl2, 0.225 % of NaHCO3, and incubated 3 days at 37°C before plaque counting.

### Infection assays

Vero cells were plated (50,000 cell per well) in p24-well plates 4 hours before being incubated with SARS-CoV-2 diluted in DMEM-2%. Freshly purified pDC were seeded in p96-well plates (50,000 cells per well) and incubated with SARS-CoV-2 diluted in pDCs culture medium (RPMI 1640 Medium with GlutaMAX, 10% of FBS, 1% of MEM NEAA, 1% of Sodium Pyruvate, and 1% of Penicillin/Streptomycin). At 2, 24 and 48 hour post-inoculation, Vero cells were trypsinized and transferred to p96-well plates. Vero and pDC were washed with PBS and fixed with 2% (v/v) paraformaldehyde diluted in PBS for 15 min at room temperature. Cells were incubated for 1 hour at 4°C with 1μg/ml of anti-nucleoprotein mAb (40143-MM05; Sino Biological) diluted in permeabilization flow cytometry buffer (PBS supplemented with 5% FBS, 0.5% (w/v) saponin, 0.1% Sodium azide). After washing, cells were incubated with 1μg/ml of Alexa Fluor 488 (115-545-003; Jackson ImmunoResearch) or 647-conjugated (115-606-062; Jackson ImmunoResearch) goat anti-mouse IgG diluted in permeabilization flow cytometry buffer for 30 min at 4°C. SARS-CoV-2 infection was quantified by flow cytometry.

To quantify infectious viral particle released in the supernatants of infected cells, pDC and Vero cells were inoculated with SARS-CoV-2 as described above and incubated at 37°C for 72-hour. At indicated time points, supernatants were collected and kept at −80°C. Virus titer was then determined by plaque assay on Vero E6 cells as described above.

### Kinetic of infection by qPCR assay

Vero E6 and pDC were inoculated with SARS-CoV-2 as described above. At the indicated time points, cells were washed thrice with PBS. Vero E6 were further incubated with trypsin 0.25% for 5 min at 37°C to remove cells surface bound particles. Total RNA was extracted using the RNeasy plus mini kit (Qiagen) according to manufacturer’s instruction. cDNAs were generated from 80 ng total RNA by using the Maxima First Strand Synthesis Kit following manufacturer’s instruction (Thermo Fisher Scientific). Amplification products were incubated with 1 Unit of RNAse H for 20 min at 37 °C, followed by 10 min at 72°C for enzyme inactivation, and diluted 10-fold in DNAse/RNAse free water. Real time quantitative PCR was performed using a Power Syber green PCR master Mix (Fisher Thermo Scientific) on a Light Cycler 480 (Roche). The primers used for qPCR were: E_Sarbeco_F1 (5’-ACAGGTACGTTAATAGTTAATAGCGT-3’), E_Sarbeco_R2 (5’-ATATTGCAGCAGTACGCACACA-3’) for viral RNA quantification. The plasmid containing the sequence corresponding to the amplified cDNA was purchased from GenScript (pUC57-2019-nCoV-PC:E; MC_0101078) and serially diluted (294 to 2.94×109 genes copies/µl) to generate standard curves.

### PDC activation

Freshly sorted pDC were cultured in p96-well plates at a concentration of 5.10^5^ cells per ml in the presence of medium alone (RPMI 1640 Medium with GlutaMAX, 10% of FBS, 1% of MEM NEAA, 1% of Sodium Pyruvate, and 1% of Penicillin/Streptomycin), influenza virus A (Charles River, A/PR/8/34, 2µg/ml), the SARS-CoV-2 primary strain 220_95 or 211_61 at a multiplicity of infection (MOI) of 1. For intracellular cytokine staining, SARS-CoV-2 was used at a MOI of 2 to activate pDC. After 24 or 48h of culture, pDC supernatant was collected and store at −80°C for subsequent cytokine quantification. PDC were stain for flow cytometry analysis.

### Flow cytometry analysis

To sort pDC, cells were stained with zombie violet or BUV fixable viability dye (Biolegend), FITC anti-CD16 (BD, clone NKP15), FITC anti-CD14 (Miltenyi, clone TÜK4), FITC anti-CD19 (Miltenyi, clone LT19), FITC anti-CD20 (BD, clone 2H7), FITC anti-CD56 (Biolegend, clone HCD56), FITC anti-CD3 (BD, clone HIT3a), BV650 or AF700 anti-CD4 (Biolegend, clone OKT4), PE-Cy7 anti-CD11c (Biolegend, clone Bu15), APC-Vio770 anti-CD2 (Miltenyi, clone LT2), and APC anti-CD5 (BD, clone UCHT2). PDCs were gated as live, lineage– (CD16, CD14, CD19, CD20, CD56 and CD3), CD2– CD5–, and CD4+ CD11c– cells.

To detect intracellular cytokines, Brefeldin A Solution 1000X (eBioscience) was added to the wells 19h after the beginning of the activation for 5h. Cells were then fixed and permeabilized using Intracellular Fixation & Permeabilization Buffer Set (eBioscience), and stained with IFNα-V450 (BD, clone 7N4-1) and TNF-α-APC (BD, clone MAb11).

For pDC diversification and checkpoint assessment, cells were stain with zombie violet fixable viability dye (Biolegend), BV711 anti-CD123 (Biolegend, clone 6H6), PE anti-CD80 (BD, clone L307.4), PerCP-efluor 710 anti-PD-L1 (eBioscience, clone MIH1), BUV737 anti-CD86 (BD, clone 2331), BV421 anti-OX40 Ligand (BD, clone ik-1), APC anti-CD62L (BD, clone DREG-56), FITC anti-CCR7 (R&D System, clone 150503).

For ACE2 cell surface expression, indicated cells were incubated with a goat anti-human ACE2 polyclonal Ab (5 μg/ml; AF933 Biotechne) in 100 μl of PBS with 0.02% NaN3 and 5% FBS for 1 h at 4°C. Cells were then washed and incubated with an Alexa 647-conjugated secondary antibody (Jackson ImmunoResearch) for 30 min at 4°C.

Acquisition was performed on an Attune NxT Flow Cytometer (Thermo Fisher Scientific) or a LSR Fortessa (BD Biosciences), and analysis was done by using FlowJo software (Tree Star). Flow cytometry analyses were performed at flow cytometry core facility of IRSL (Paris, France).

### Secreted inflammatory cytokines measurement

PDC cytokine production of IFN-α2, IL-8, IL-6, IP-10 and TNF-α, was measured in culture supernatants using BD cytometric bead array (CBA), according to the manufacturer’s protocol, with a 20pg/ml detection limit. Acquisitions were performed on a LSR Fortessa (BD Biosciences), and cytokine concentrations were determined using FCAP Array Software (BD Biosciences).

The concentration of secreted IFN-λ1 was measured by enzyme-linked immunosorbent assay (ELISA) (R&D Systems, DuoSet DY7246), according to the manufacturer’s instructions. The manufacturer reported neither cross-reactivity nor interference with IFN-α, IFN-β 1a, IL-10Rβ, IFN-λ2 and λ3, and IL-28Rα. The optical density value (OD) of the supernatants was defined as its absolute OD value, minus the OD Absorbance from blank wells. The detection limit was 85pg/ml and all samples were run in duplicates.

### Statistical analysis

Statistical analyses were performed with one-way ANOVA, Kruskall Wallis’s test with Dunn’s multiple comparison post-test or Mann Whitney’s test, in Prism (GraphPad Software).

## Online supplemental material

Fig. S1 A, B and C can relate to all of this paper. Fig. S1 A, B presents that magnetically sorted, frozen pDC diversification is similar to FACS, fresh pDC when activated with SARS-CoV-2 or Flu, demonstrating that the technical limitations we encounter during this work did not impact its relevance. Data shown in Fig. S1 C, highlights pDC diversification when activated with either free SARS-CoV-2, or SARS-CoV-2 infected cells. Fig. S1 D and E gives additional evidence from Fig. 1 that pDC are not permissive to SARS-CoV-2 infection. Fig. S2 provides additional information on Fig. 1, 2 and 3. Fig. S2 A, B and C show pDC diversification and pro-inflammatory cytokines after 48h of culture with either Medium, SARS-CoV-2 or Flu. Fig S2. D and E demonstrate that tonsillar pDC can also undergo diversification and pro-inflammatory cytokine secretion when activated with Flu or SARS-CoV-2 compared to Medium. Fig S3 relates to Fig 4. Fig S3 A shows HCQ potential to inhibit pDC diversification in a dose-dependent manner. Fig S3 B and C demonstrate the absence of OX40L^high^ pDC when cultured with HCQ, providing further evidence that HCQ has inhibitory effects on pDC diversification.

## Author contributions

Ali Amara and Vassili Soumelis conceived the study. Fanny Onodi, Lucie Bonnet-Madin, Ali Amara and Vassili Soumelis designed the experiments. Fanny Onodi performed the pDC purification and FACS analysis with Justine Poirot. Lucie Bonnet-Madin and Laurent Meertens performed the virus strain isolation and the infection studies. Fanny Onodi and Léa Karpf performed pDC purification and cytokine quantification. Jean-Michel Molina, Constance Delaugerre and Jérome Le Goff performed the sequencing of the SARS-CoV-2 strains. Shen-Ying Zhang, Anne Puel, Emmanuelle Jouanguy, Qian Zhang and Jean-Laurent Casanova coordinated recruitment and provided patient samples. Jean-Laurent Casanova revised the manuscript. Capucine Picard provided patient samples and collected clinical data. Fanny Onodi, Ali Amara and Vassili Soumelis wrote the initial manuscript draft and all authors contributed to the final form of the manuscript.

## Acknowledgments

Ali Amara’s lab received fundings from the French Government’s Investissement d’Avenir program, Laboratoire d’Excellence “Integrative Biology of Emerging Infectious Diseases” (grant n°ANR-10-LABX-62-IBEID), the Fondation pour la Recherche Medicale (grant FRM - EQU202003010193), the Agence Nationale de la Recherche (ANR-FRM FLASH COVID project IDISCOVR), University of Paris (Plan de Soutien Covid-19: RACPL20FIR01-COVID-SOUL). Vassili Soumelis’ team was supported by Agence Nationale de la Recherche (ANR-17-CE15-0003, ANR-17-CE15-0003-01), and by Université de Paris « PLAN D’URGENCE COVID19 ». The Laboratory of Human Genetics of Infectious Diseases is supported by the Howard Hughes Medical Institute, the Rockefeller University, the St. Giles Foundation, *Institut National de la Santé et de la Recherche Médicale (*INSERM), *Université de Paris*, the French Foundation for Medical Research (FRM) (EQU201903007798), the French National Research Agency (ANR) under the “Investments for the Future” program (ANR-10-IAHU-01), the Integrative Biology of Emerging Infectious Diseases Laboratory of Excellence (ANR-10-LABX-62-IBEID), the CNSVIRGEN project (ANR-18-CE15-0009-01) and the SEAe-HostFactors project (ANR-18-CE15-0020-02). the National Institutes of Health (NIH) (R01AI088364), the National Center for Advancing Translational Sciences (NCATS), NIH Clinical and Translational Science Award (CTSA) program (UL1 TR001866), a Fast Grant from Emergent Ventures, Mercatus Center at George Mason University, the Yale Center for Mendelian Genomics and the GSP Coordinating Center funded by the National Human Genome Research Institute (NHGRI) (UM1HG006504 and U24HG008956), the FRM and ANR GENCOVID project, ANRS-COV05, the Square Foundation, the Fisher Center for Alzheimer’s Research Foundation and *Grandir - Fonds de solidarité pour l’enfance*. The authors thank Maud Salmona, Severine Mercier-Delarue and Tiffanie Bouillé (Laboratoire de Virologie, Hôpital Saint Louis) for SARS-CoV-2 deep sequencing, Marie Laure Chaix (Laboratoire de Virologie, Hôpital Saint Louis) for the nasopharyngeal swabs and Lauriane Goldwirt (Laboratoire de Pharmacologie Biologique, Hôpital Saint-Louis) for providing us with hydroxychloroquine. We warmly thank S. Boucherit for administrative assistance. The authors thank all the “Personnel Soignant de l’Hôpital Saint-Louis” for their amazing work during the COVID-19 epidemic. Ali Amara dedicates this work to the memory of Pr Jean-Louis Virelizier (Unité d’Immunologie Virale, Institut Pasteur, Paris) and Pr Renaud Mahieux (ENS and CIRI, Lyon, France), who left us during the pandemic.

## Abbreviations

CBA: Cytometric bead array
COVID-19: Coronavirus disease 2019
CPE: Cytopathogenic effects
DC: Dendritic cell
Flu: Influenza virus A
HCQ: Hydroxychloroquine
HD: Healthy donor
IFN: Interferon
MERS: Middle East respiratory syndrome coronavirus
MFI: Mean fluorescence intensity
MHV: Murine hepatitis virus
MOI: Multiplicity of infection
N: nucleoprotein antigen
OD: optical density
PBMC: Peripheral blood mononuclear cells
pDC: Plasmacytoid pre-dendritic cells
SARS-CoV-2: Severe Acute Respiratory Syndrome-Coronavirus-2
TLR: Toll-like receptor

---

**Figure S1. SARS-CoV-2 induces pDC activation**. Sorted blood pDC from healthy donors were cultured for 24h with either Medium, SARS-CoV-2, or Influenza virus A (Flu). **(A)** Percentage of pure pDC among live cells through different sorting strategies. Results from two healthy donors representative of n=8. **(B)** P1-, P2- and P3-diversification of fresh, fluorescent sorted pDC versus frozen, magnetic sorted pDC with either SARS-CoV-2 or Influenza virus A (Flu) for 24h. Results from two healthy donors representative of n=8. **(C)** Dotplot of pDC activation with either free SARS-CoV-2 or pDC co-culture with SARS-CoV-2 infected cells. Results from n=1 healthy donors. **(D)** Viral RNA copy number in of Vero E6 and pDC 2, 24 and 48h post infection (hpi). Results from one experiment representative of n=3. **(E)** Intracellular production of the nucleoprotein antigen (N) on pDC of two healthy donors of n=3.

**Figure S2. SARS-CoV-2 induces activation and diversification of tonsilar pDCs**. Sorted blood pDC from healthy donors were cultured for 48h with Medium, SARS-CoV-2, or Influenza virus A (Flu). **(A)** Dotplot showing pDC activation and diversification through the expression of PD-L1 and CD80 at 24h and 48h. Results from one healthy donors representative of n=3. **(B)** Geometric mean (MFI) of pDC’s activation markers at 48h. Histograms represent medians and bars interquartiles of n=3 healthy donors from 3 independent experiments. *P□< 0.05; Kruskal-Wallis with Dunn’s multiple comparison’s post test. **(C)** Quantification of secreted pro-inflammatory cytokines 48h after culture. Bars represent medians of n=3 healthy donors from 3 independent experiments. *P□< 0.05; ns: not significant; Mann Whitney test. **(D)** Dotplot of tonsil pDC activation cultured for 24h with either Medium, SARS-CoV-2 or Influenza virus A (Flu). Results from n=1 healthy donors. **(E)** Quantification of pro-inflammatory cytokines of tonsilar pDC at 24h. Histograms represent medians of n=1 healthy donor. ND: Not Detectable.

**Figure S3. Hydroxychloroquine inhibits SARS-CoV-2-induced pDC activation in a dose dependent manner**. Sorted blood pDC from healthy donors were cultured for 24h with either Medium, Influenza virus A (Flu), or SARS-CoV-2 at a MOI 1 with hydroxychloroquine (HCQ) or vehicule. **(A)** Dotplot of pDC diversification with increasing concentration of hydroxychloroquine (HCQ) or vehicule. Results from one healthy donors representative of n=3. **(B)** Dotplot showing OX40L and CD86 in presence or absence of HCQ. Results from one healthy donors representative of n=3. **(C)** Percentage of OX40L^high^ population among pDC. Bars represent medians of n=3 healthy donors from 3 independent experiments. *P□< 0.05; Mann Witney test.

