## Supplementary figures and images for "SARS-CoV-2 induces human plasmacytoid pre-dendritic cell diversification via UNC93B and IRAK4"

### Supp Figure S1

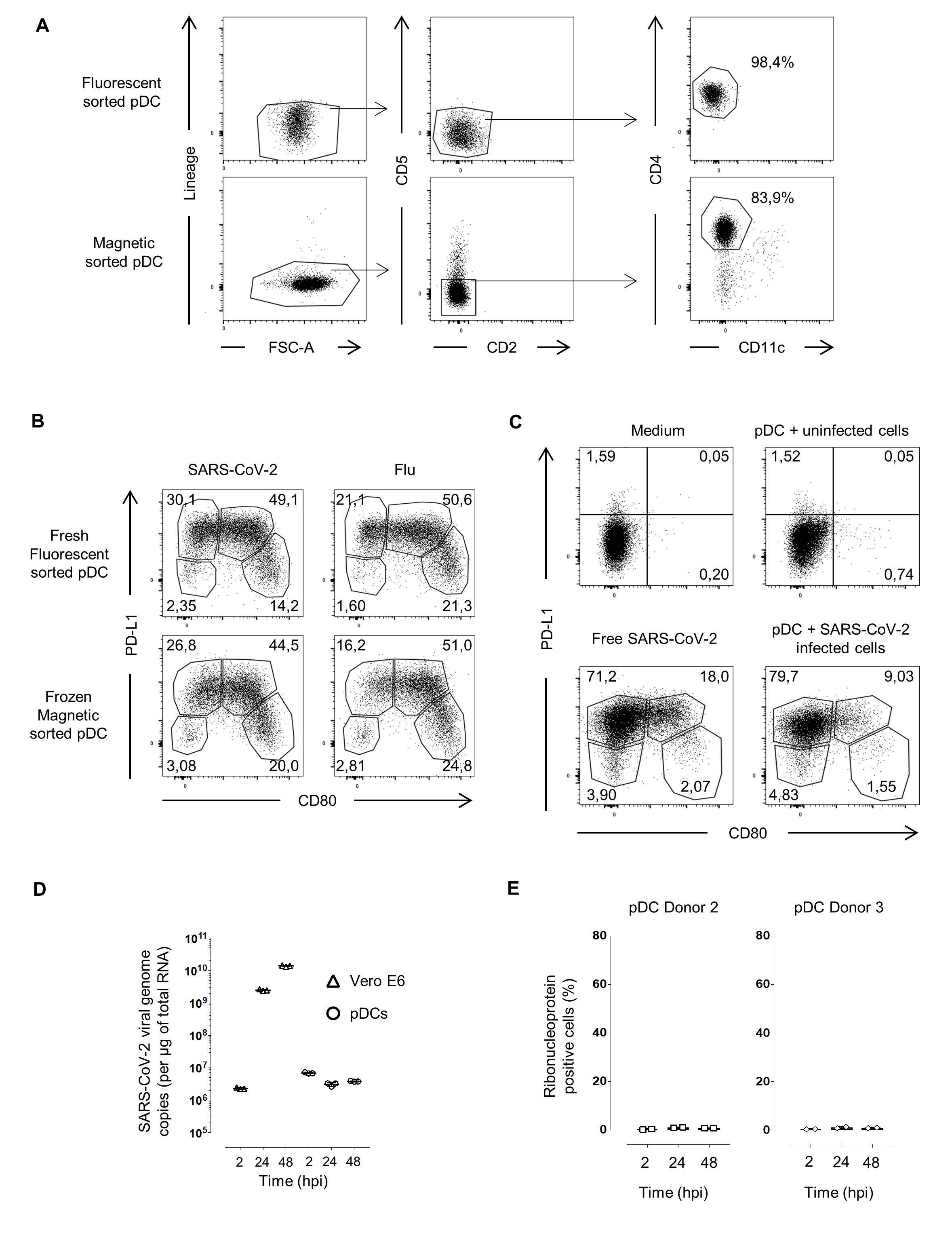

### Supp Figure S2

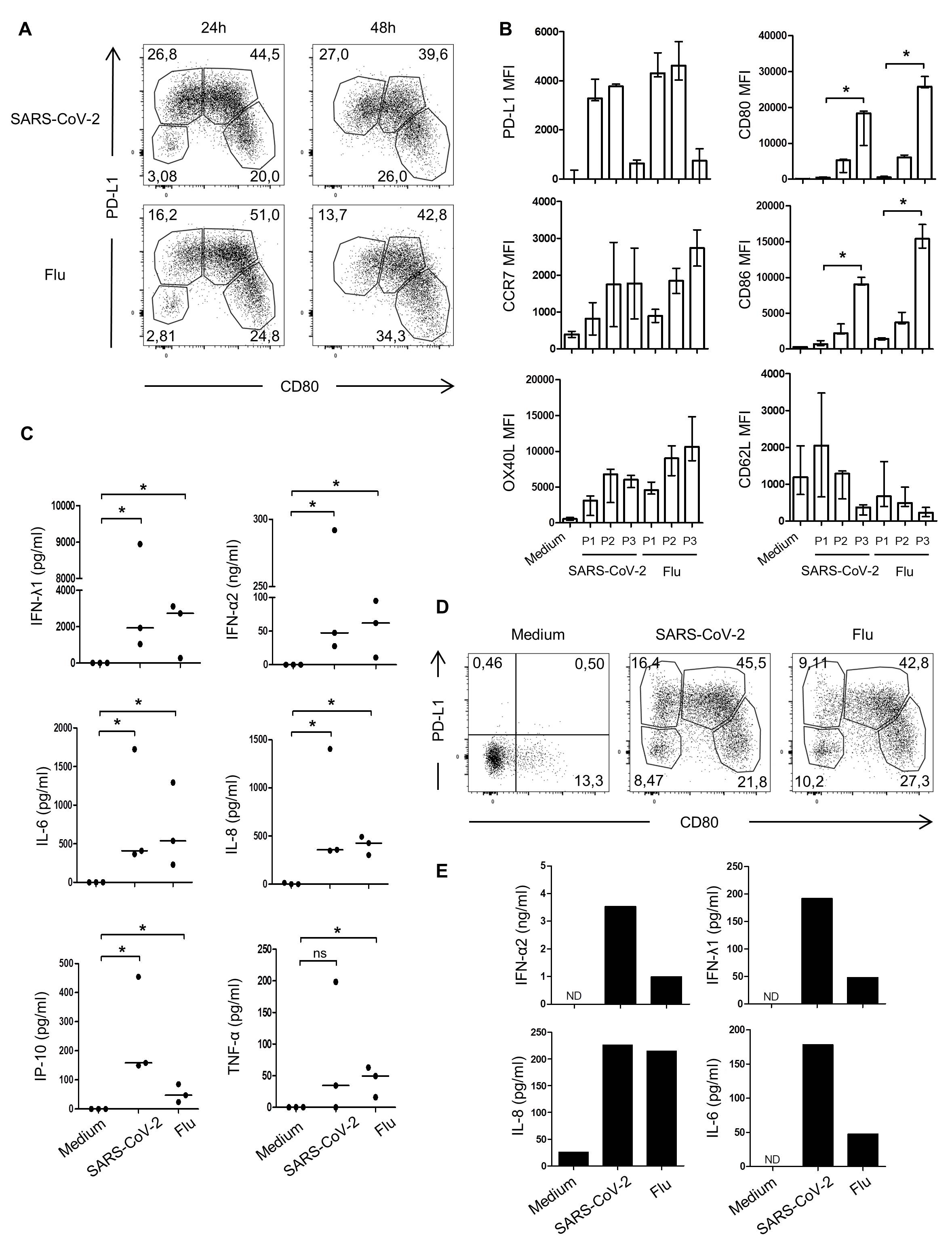

### Supp Figure S3

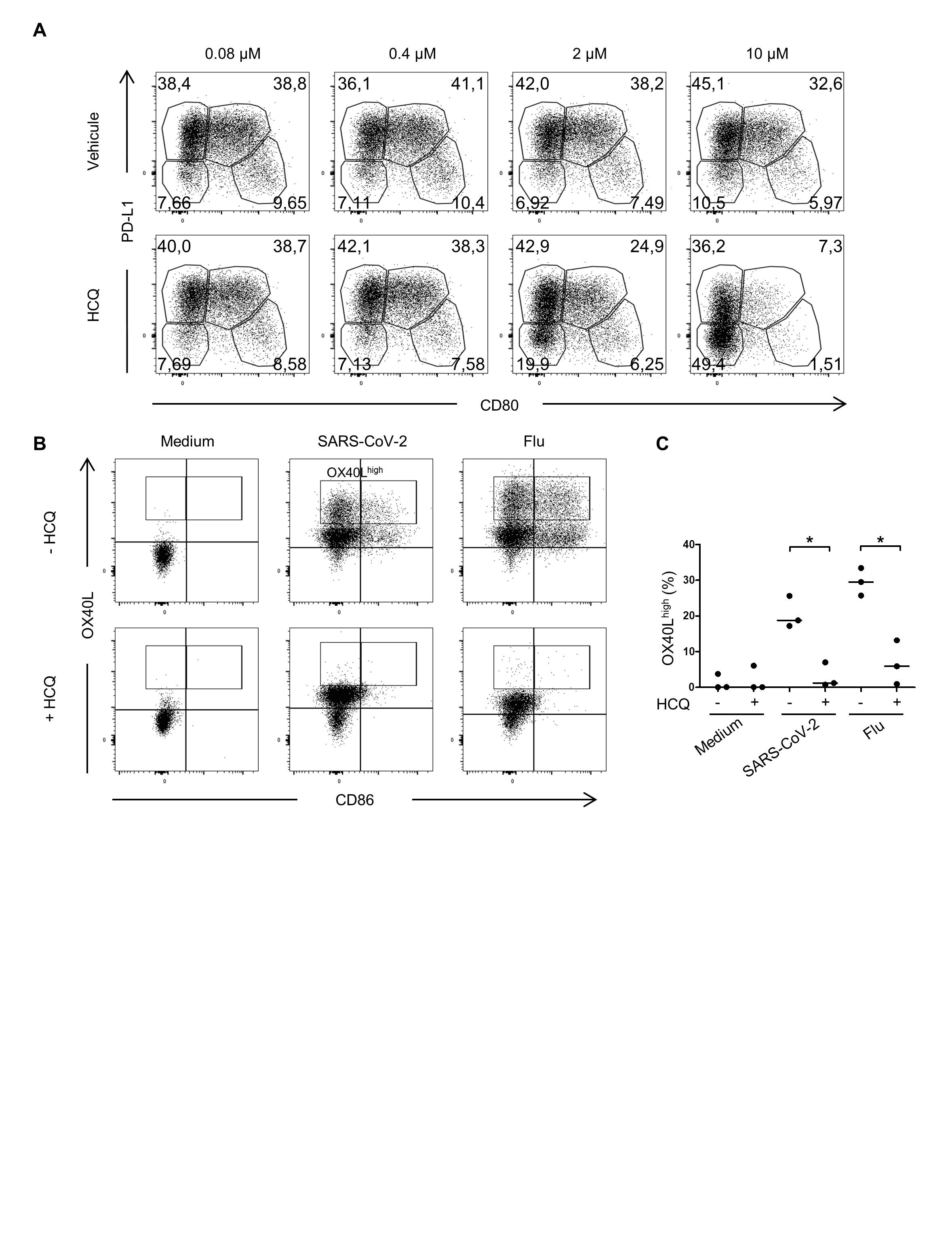
